# Quantifying Feral Pig Interactions to Inform Disease Transmission Networks

**DOI:** 10.1101/2024.08.31.610621

**Authors:** Tatiana Proboste, Abigail Turnlund, Andrew Bengsen, Matthew Gentle, Cameron Wilson, Lana Harriott, Richard A. Fuller, Darren Marshall, Ricardo J. Soares Magalhães

## Abstract

Feral pigs threaten biodiversity in 54 countries worldwide and cause an estimated $120 billion of damage annually in the United States of America (USA). Feral pigs imperil over 600 native species and have directly driven 14 species into extinction. Moreover, feral pig populations pose a significant zoonotic disease threat to humans such as Japanese encephalitis, and act as reservoir for endemic pathogens such as *Brucella* and leptospirosis. Efforts to understand and control disease spread by feral pigs rely on models of social dynamics, how the animals interact with one another. Yet social dynamics are known to vary enormously from place to place, so knowledge generated for example in USA and Europe might not easily transfer to locations such as Australia.

Here, we fill a continental gap in our understanding of feral pig social dynamics by developing a proximity-based social network analysis approach to rapidly assess social interactions using animal tracking data. This method, applied to the continent of Australia, included 146 GPS-monitored feral pigs, and revealed distinct patterns influenced by sex and season, with females demonstrating higher group cohesion (female-female) and males acting as crucial connectors between independent groups. Contact rates are remarkably high within groups, indicating rapid intra-group disease spread that contrasts with much slower potential for inter-group disease spread. Seasonal variations further complicate this dynamic, with contact rates being much higher in summer. The results show that, in Australia, targeting adult males in feral pig control programs could enhance efforts to contain disease outbreaks.

Concern over the economic and human health impacts of animal diseases is higher than ever before. We urge a rapid global effort to use models of feral pig social interactions to develop efficient control strategies tailored to local conditions.

## INTRODUCTION

Feral pigs (*Sus scrofa*) are threaten biodiversity in 54 countries worldwide [1], and cause an estimated $120 billion of damage annually in the US [2]. Feral pigs imperil over 600 native species, and have directly driven 14 species to extinction [1], and cause significant economic lost to agricultural industries through negative impacts on water quality, extensive ground rooting, lamb predation, and crop and pasture consumption [3].

Feral pigs can be reservoir hosts of multiple infectious diseases of economic and/or animal health significance. This is particularly important for the continent of Australia, where many diseases are exotic, including foot-and-mouth disease (FMD), swine vesicular disease, Aujeszky’s disease, and African and Classical swine fever (ASF and CSF). The potential economic impact of these diseases is substantial. For instance, an incursion of FMD is estimated to cost Australia $50 billion [4] and the incursion of ASF can cost up to 2.5 billion [5]. Feral pig populations in Australia can also be a source of zoonotic pathogens of public health importance including *Brucella suis* [6], *Leptospira interrogans* [7], *Coxiella burnetii* [8] and Japanese encephalitis virus [9]. Despite ongoing control measures that aim to reduce feral pig impacts, the threat of spillover of infectious diseases to domestic animals, wildlife, and human populations in Australia remains significant due to persistent feral pig populations.

### What is Australia doing to avoid the impact?

National preparedness for the incursion of diseases that are spread by feral pig activity relies on the development and validation of disease transmission models. Australia invests significant resources as part of the planning and prevention of new disease incursions and has developed the Australian Animal Disease Spread (AADIS) model [10] for ASF and FMD. However, the performance of AADIS relies on adequate parameterisation of real-world feral pig contact rates, which is particularly challenging to obtain and estimate in wild pig populations [11]. Currently, contact rate estimates needed to optimise the AADIS were obtained from European feral pig population studies [12-14], and remain unavailable for Australian feral pig populations. Locally relevant data on feral pigs contact rates will provide more reliable parameters to model the disease transmission rate in feral pig populations.

### Contact rates are imperative for disease models

To estimate disease spread, the force of infection is needed (β), which can be estimated by knowing the population of the contact rate (γ), the probability that contact is made with infected individuals and the probability of pathogen transmission given a contact (*K*) [11, 15]. Using individual movement network data, we can estimate the contact between individuals in wild populations. Contact rates between individual feral pigs are affected by feral pig social structure and spatial distribution [16]. Historically, estimating contact rates from GPS tracking data of individuals has been challenging due to the need to set arbitrary time or distance thresholds to define a ‘contact’ or from limitations in the interval of the GPS collar recording location. New approaches have been recently developed to refine these calculations and to account for spatial autocorrelation, like continuous-time movement modelling (CTMM), which downscales the recording interval and improves the detection of contacts between individuals [17].

Understanding how animals move in space and time is critical to estimating the rate of infectious disease transmission and to optimising disease transmission models [18], to improve the confidence in projected impacts of modelled population control measures. Currently, there is a lack of understanding of contact rates between and within feral pigs, information that is necessary for the development of exotic disease model and contingency plans [19]. Therefore, it is vital to examine feral pig movement behaviours in a heterogeneous landscape, contact rates between individuals, differences in contact rates within and between pig social units (sounders), and to identify sex and age classes which are the most likely to transmit infections in case of an outbreak.

Contact heterogeneities for feral swine or wild boar have been estimated in different regions. In Poland, significant clustering of individuals was detected and differences in the duration of these associations depended on the sex of the individual [20]. This reaffirms that dominant boars are primarily solitary, while reproductively active sows and their offspring largely remain in sounders [21, 22]. Feral pig movement and interactions have also been explored with network analysis to estimate contact rates of populations in the USA [23], Germany and Italy [14]. For example, the US study found the contact rate between sounders was affected by the distance between the home range, with less contact when home ranges were separated by more than 2km [23]. The European study found similar results and described that the most frequent association occurred at distances of 0-1 km and more sporadic association at more than 4km [14]. However, such data have not been analysed for feral pigs in Australia.

This study aims to fill a continental gap in our understanding of feral pig social dynamics within and between sounders in eastern Australia, and to investigate differences between direct and indirect contacts. Our goal is to identify specific characteristics, such as demographic variables, which are common among individuals who are central to these networks. The findings from this research will enhance existing ASF models, as well as other pig disease transmission models, by providing empirical data on the structure of feral pig contact networks. Importantly, this approach allows us to identify key individuals that are central to these networks and, therefore, more likely to facilitate the spread of pathogens. This crucial information will greatly enhance the precision and relevance of disease modelling within feral pig populations to inform persistence, spread and the design and implementation of management actions.

## Results

A total of 139,940 location fixes were collated from 146 GPS-monitored feral pigs, 61 females and 85 males, tracked from 2017 to 2023 across QLD and NSW (Figure 1). Arcadia (QLD) population provided the most fixes (403,313) and the largest sample size (n = 31), while Nap Nap (NSW) provided the least fixes (18,637) and had the smallest sample size (n = 2). We obtained the most fixes during spring with 27%, 25% fixes during winter, and 24% fixes during autumn and 24% fixes during summer season from 2017 to 2023 across QLD and NSW.

**Figure 1.**
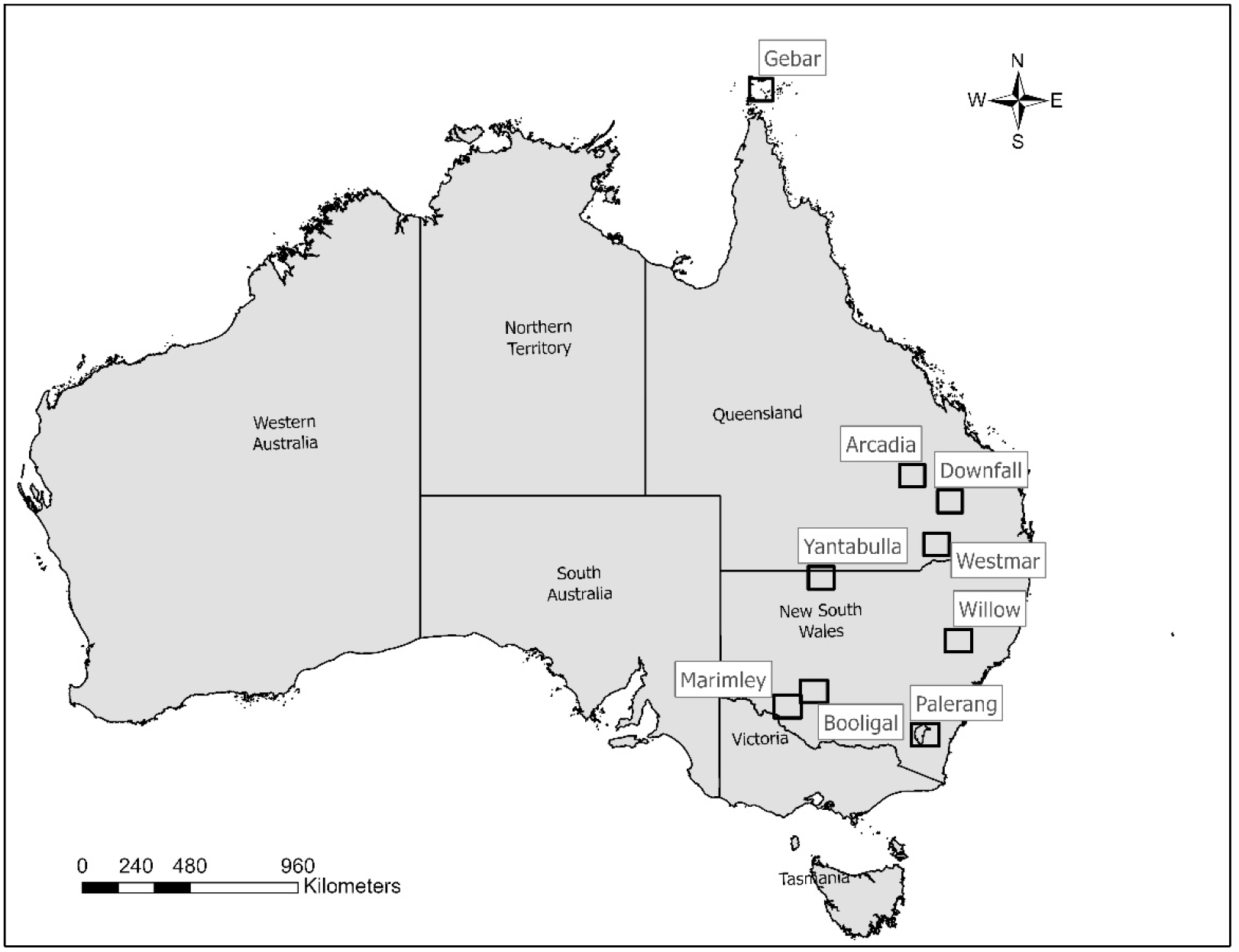
Map of Australia identifying the location (name of the population) of the study site.

We compared the global network measures derived from thresholds of 2, 5, and 350 metres (Figures S1-2). The results revealed differences between the 2 and 5-metre thresholds in terms of average local transitivity, with an increase in the number of clusters as we raised the thresholds. Regarding edge density, which provides insight into the level of interconnectivity within the networks, our findings showed similar patterns across the thresholds, with a trend of increased interconnectivity as the threshold rose. The mean distance, which calculates the length of the shortest path for all possible pairs in the networks, showed minimal differences across the various thresholds.

The comparison of the global network measures between the direct and indirect networks (5 metres threshold), revealed that average local transitivity, edge density and global transitivity were slightly higher for the indirect networks than the direct network (Figure S3). Pigs within the Yantabulla population had a higher betweenness as well as a higher degree than pigs at other sites. Additionally, betweenness in males was, on average, 1.62 times greater than in females (Figure 2).

**Figure 2.**
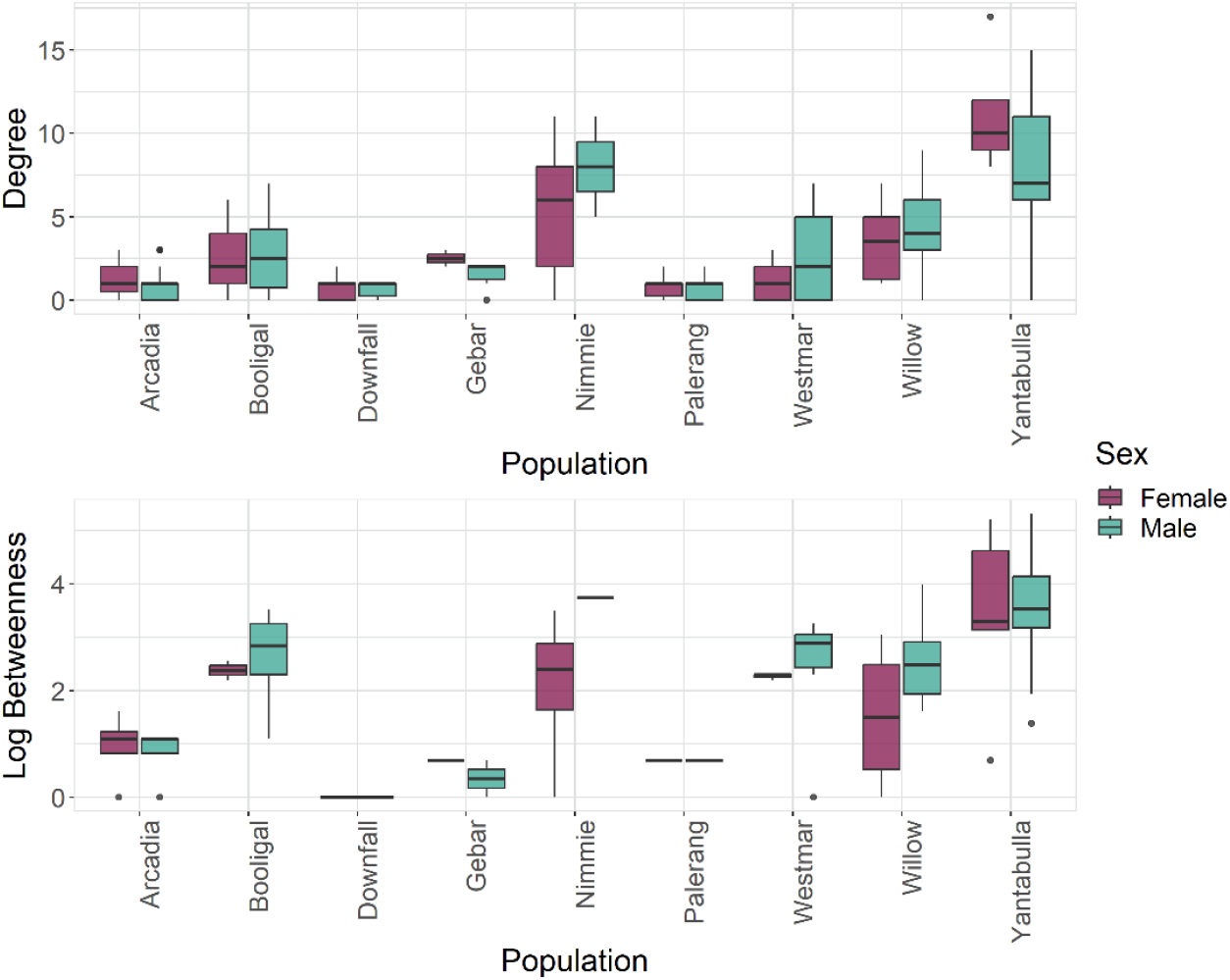
Betweenness (log) and degree measures at the individual level for each population and by sex, where green represent males and red represents females.

Comparison of node-level network measures between females and males for the direct network and indirect network based on a 5 metres threshold (Figure 3) revealed that males were positively associated with betweenness (log) (statistical significant for indirect network), while females were positively associated with higher strength (statistical significant for direct network) (Table 1).

**Table 1.**
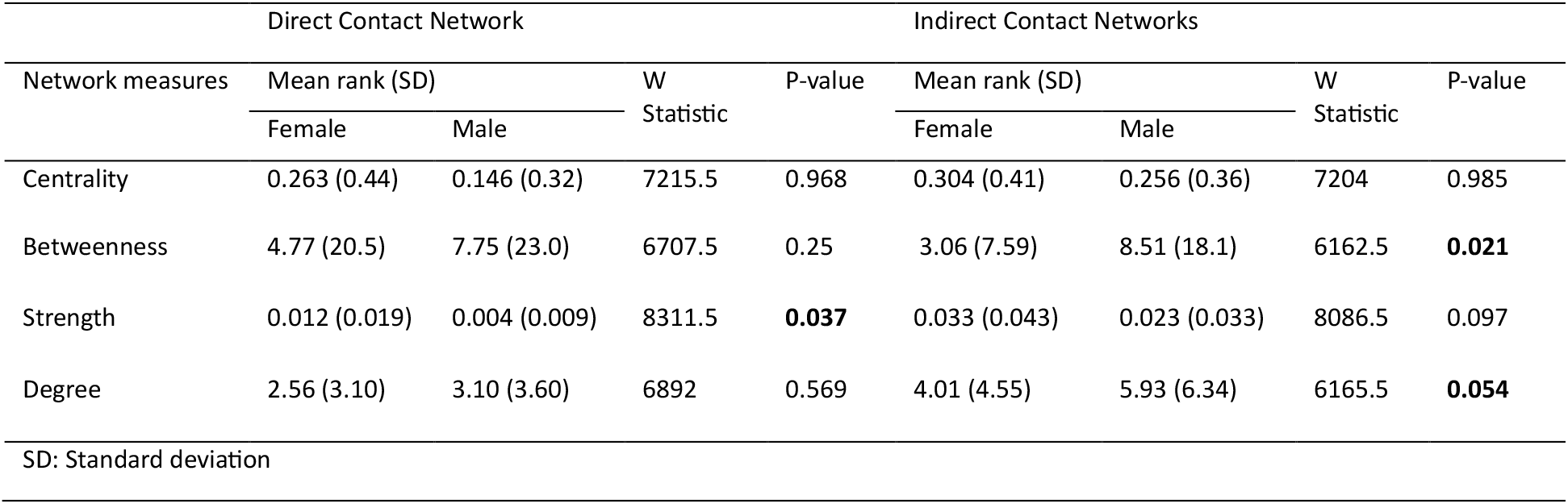
Differences between sex and network measures using Wilcoxon rank-sum test for direct and indirect contact.

| Network measures | Direct Contact Network |  |  |  | Indirect Contact Networks |  |  |  |
| --- | --- | --- | --- | --- | --- | --- | --- | --- |
|  | Mean rank (SD) |  | W<br>Statistic | P-value | Mean rank (SD) |  | W<br>Statistic | P-value |
|  | Female | Male |  |  | Female | Male |  |  |
| Centrality | 0.263 (0.44) | 0.146 (0.32) | 7215.5 | 0.968 | 0.304 (0.41) | 0.256 (0.36) | 7204 | 0.985 |
| Betweenness | 4.77 (20.5) | 7.75 (23.0) | 6707.5 | 0.25 | 3.06 (7.59) | 8.51 (18.1) | 6162.5 | **0.021** |
| Strength | 0.012 (0.019) | 0.004 (0.009) | 8311.5 | **0.037** | 0.033 (0.043) | 0.023 (0.033) | 8086.5 | 0.097 |
| Degree | 2.56 (3.10) | 3.10 (3.60) | 6892 | 0.569 | 4.01 (4.55) | 5.93 (6.34) | 6165.5 | **0.054** |
| SD: Standard deviation |  |  |  |  |  |  |  |  |

**Figure 3.**
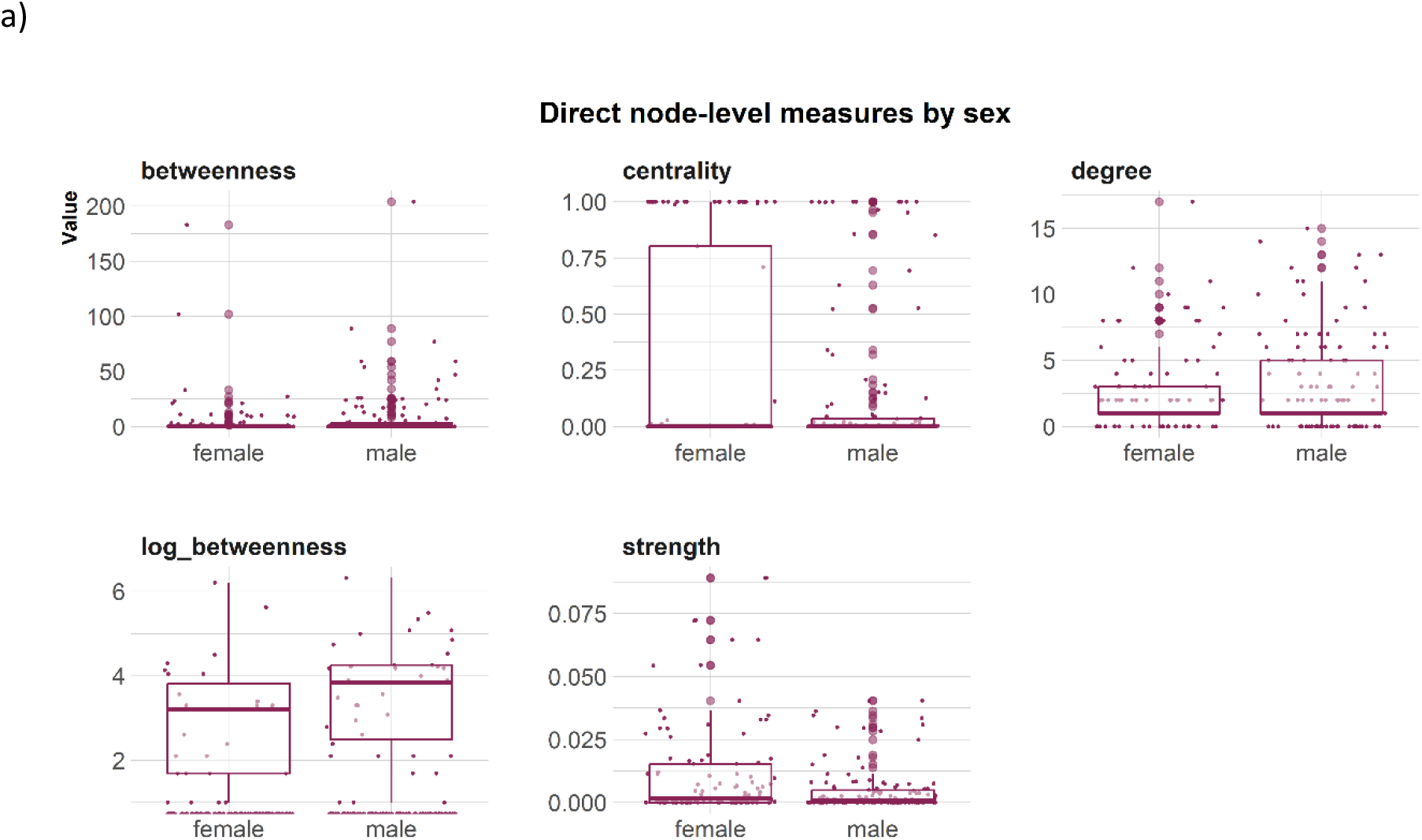
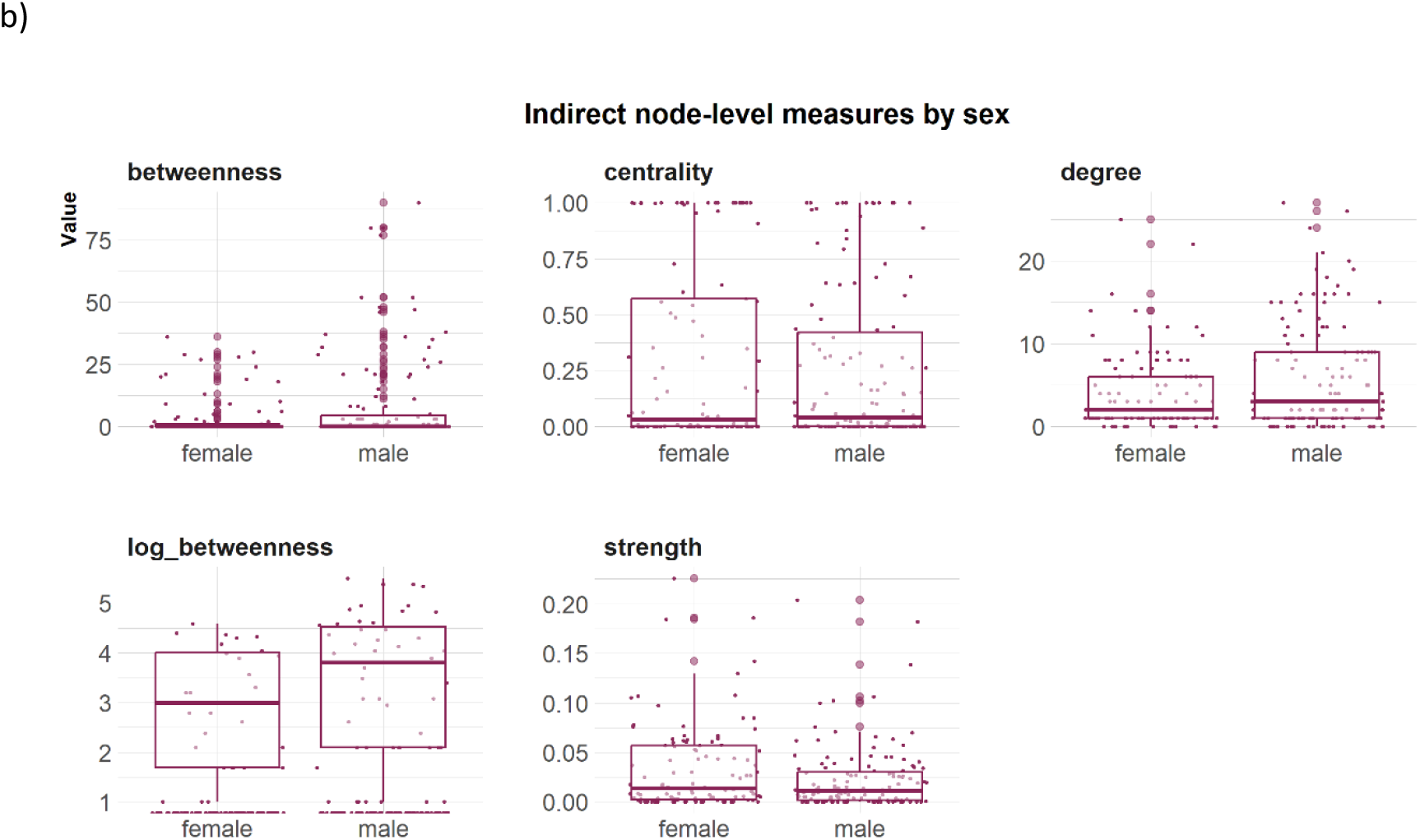
Node-level measures (5m threshold), including betweenness, centrality, degree, log(betweenness) and strength by sex for a) direct network and b) indirect network.

### Home range

Home range overlap within dyads was greatest during summer, followed by autumn; however, this difference was not statistically significant (Figure 4).

**Figure 4.**
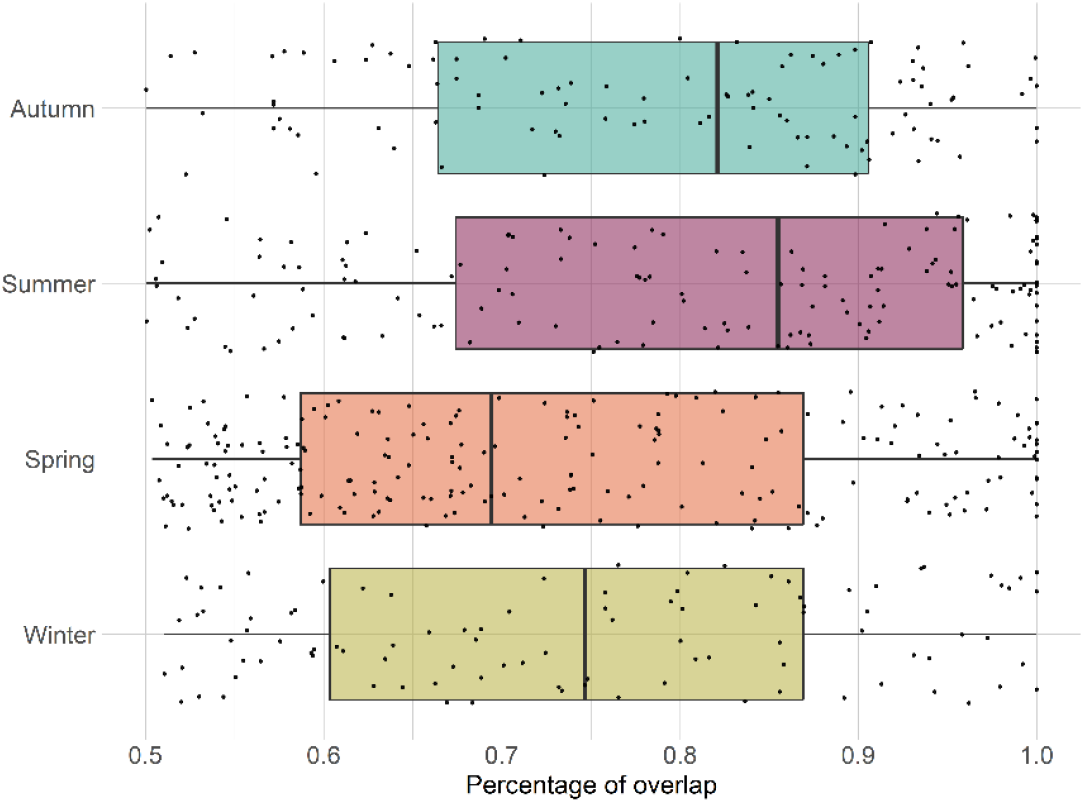
Home range overlap per dyad for each season.

### Contact rates

Most of the records of direct (96%) and indirect contact (69%) occurred between animals from the same sounder. Most observations of indirect contact within and between the sounders occurred during winter and the least number of observations occurred during summer. We found a positive effect (p-values <0.001) on the contact rate within sounders compared to contact between sounders (overall contact rate in Table 2 and Figure 5) and that the contact rate differed depending on the sex of the dyad.

**Table 2.**
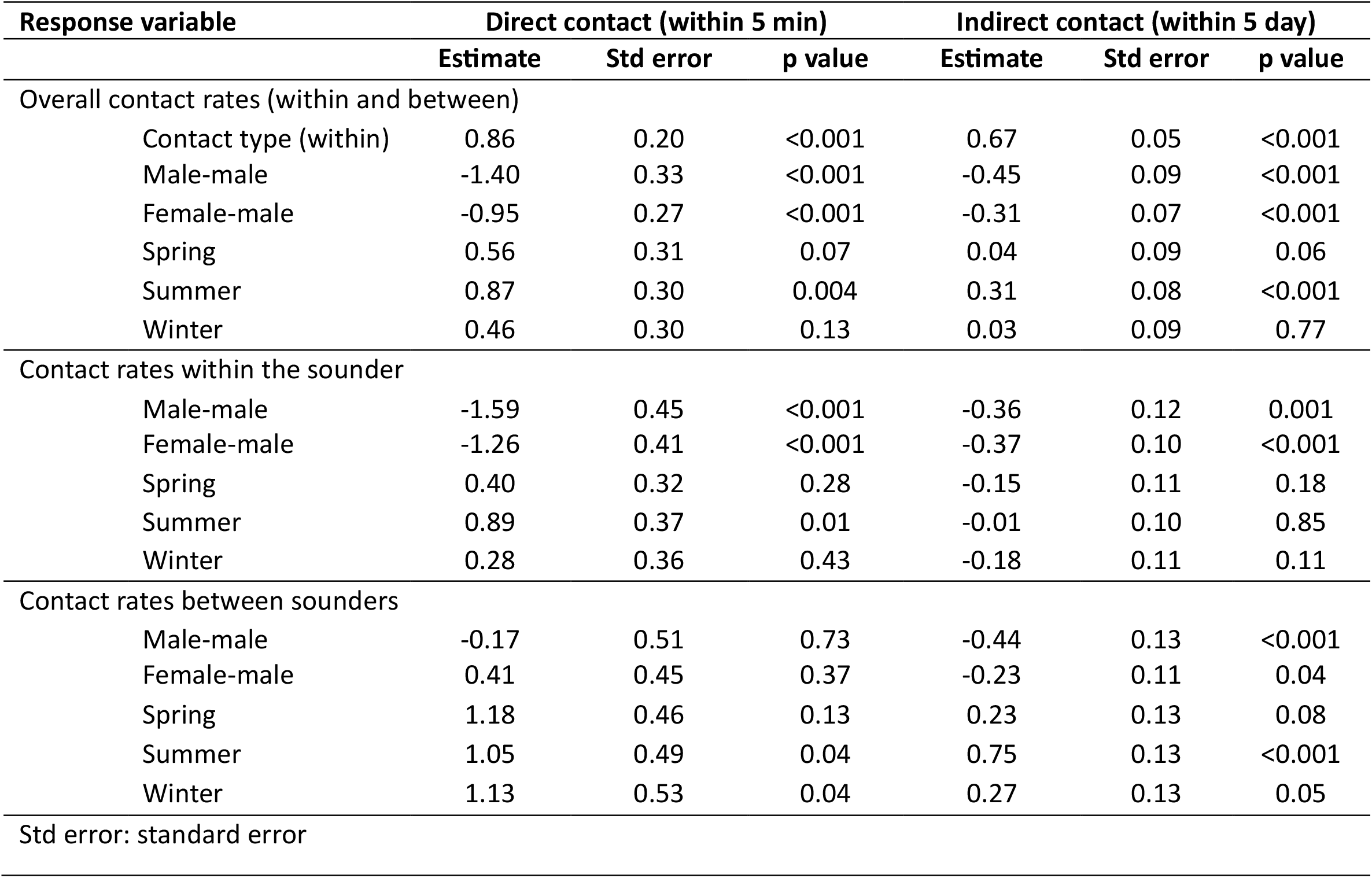
Association between the mean contact rate within sounders, between sounders and overall and the distinct types of contacts (female-female, male-male, female-male) and seasons (autumn, spring, summer, winter), for direct and indirect contact calculated with 5 metres threshold.

| Response variable | Direct contact (within 5 min) |  |  | Indirect contact (within 5 day) |  |  |
| --- | --- | --- | --- | --- | --- | --- |
|  | Estimate | Std error | p value | Estimate | Std error | p value |
| Overall contact rates (within and between) |  |  |  |  |  |  |
| Contact type (within) | 0.86 | 0.20 | <0.001 | 0.67 | 0.05 | <0.001 |
| Male-male | -1.40 | 0.33 | <0.001 | -0.45 | 0.09 | <0.001 |
| Female-male | -0.95 | 0.27 | <0.001 | -0.31 | 0.07 | <0.001 |
| Spring | 0.56 | 0.31 | 0.07 | 0.04 | 0.09 | 0.06 |
| Summer | 0.87 | 0.30 | 0.004 | 0.31 | 0.08 | <0.001 |
| Winter | 0.46 | 0.30 | 0.13 | 0.03 | 0.09 | 0.77 |
| Contact rates within the sounder |  |  |  |  |  |  |
| Male-male | -1.59 | 0.45 | <0.001 | -0.36 | 0.12 | 0.001 |
| Female-male | -1.26 | 0.41 | <0.001 | -0.37 | 0.10 | <0.001 |
| Spring | 0.40 | 0.32 | 0.28 | -0.15 | 0.11 | 0.18 |
| Summer | 0.89 | 0.37 | 0.01 | -0.01 | 0.10 | 0.85 |
| Winter | 0.28 | 0.36 | 0.43 | -0.18 | 0.11 | 0.11 |
| Contact rates between sounders |  |  |  |  |  |  |
| Male-male | -0.17 | 0.51 | 0.73 | -0.44 | 0.13 | <0.001 |
| Female-male | 0.41 | 0.45 | 0.37 | -0.23 | 0.11 | 0.04 |
| Spring | 1.18 | 0.46 | 0.13 | 0.23 | 0.13 | 0.08 |
| Summer | 1.05 | 0.49 | 0.04 | 0.75 | 0.13 | <0.001 |
| Winter | 1.13 | 0.53 | 0.04 | 0.27 | 0.13 | 0.05 |
| Std error: standard error |  |  |  |  |  |  |

**Figure 5.**
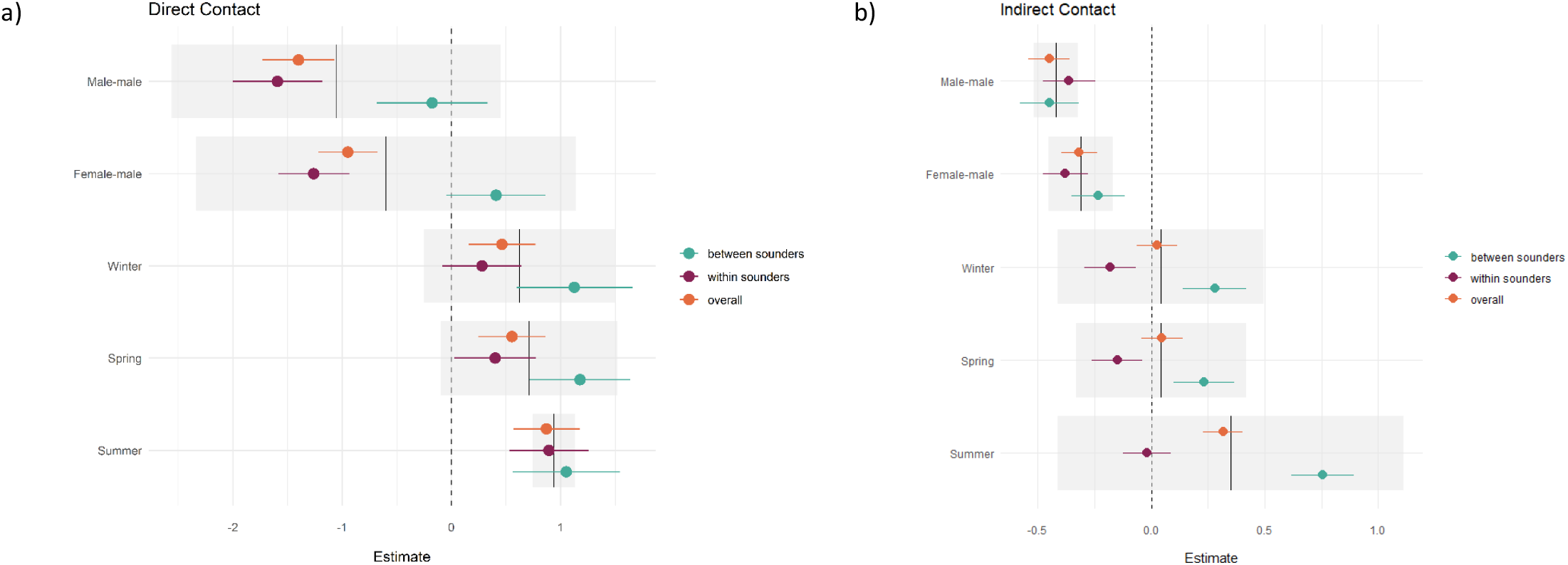
Distribution of the estimates and the effect of dyad sex and season in the contact rate a) during direct contact and b) for indirect contact. References are female-female for the sex of the dyad and autumn for the season.

Direct contact rates for female-female dyads across all interactions (within and between sounders) were 1.4 times greater than those for male-male dyads (estimate =-1.4; p <0.001) and 0.95 greater than those for male-female dyads (estimate = -0.95, p <0.001; Figure 5a).

Within sounders, we observed a similar trend, where female-female dyad were 1.56 times greater than male-male and 1.2 times greater than female-male, and contacts were more frequent during summer. We did not find a significant effect of sex of the dyad for contact rate between sounders, and contacts were more frequent during summer and winter compared to autumn.

For the indirect overall contacts (Figure 5b), we found that male-male and female-male contact were less frequent than female-female contact (p-values <0.001), and the contact rate was more frequent in summer compared to autumn (p-value <0.001). For the contact rate within sounders, we found a similar trend, with less frequent contact between female-male and male-male. Between sounders, male-male contact was less frequent compared to female-female contact, and summer had the highest contact frequency (p-value <0.001; Table 2).

Mean contact between sounders was generally lower than the mean contact within sounders. Within sounders, the highest mean contact occurred between dyads of females, and it was also higher for Arcadia (QLD) and Booligal (NSW) populations; both cases population the contact rate increased with the years (Supplementary material). No contact was detected within sounders for two populations, Marimley and Nap Nap, as these populations had very small sample sizes (Figure 6).

**Figure 6.**
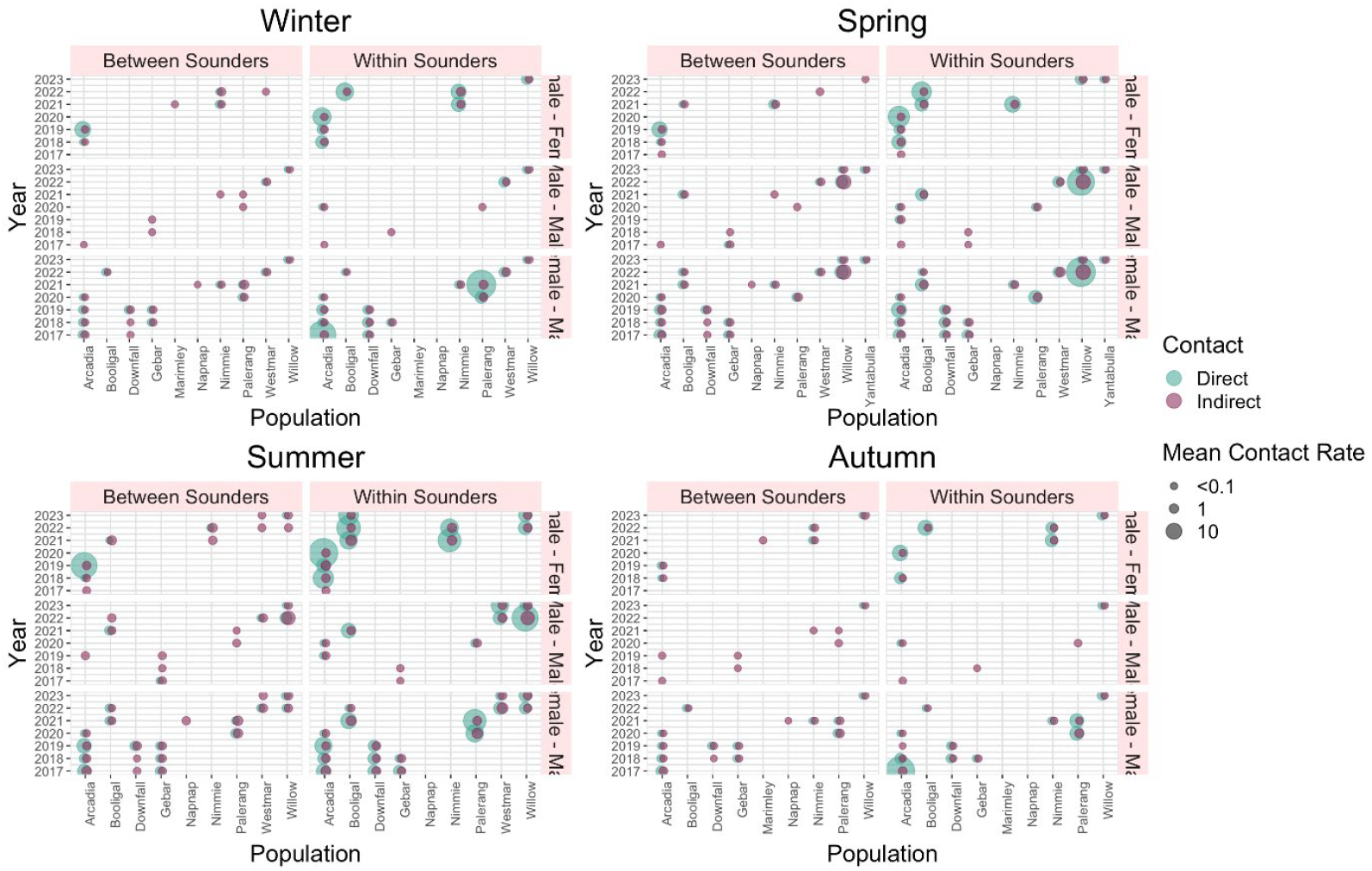
Diagram of direct and indirect mean contact rate for each season for dyads between sounders and within sounders per year and population

## Discussion

Models of disease spread provide important insights for preparedness and response. The incursion of an exotic enzootic disease such as African Swine Fever (ASF) or Foot-and-mouth Disease (FMD) could result in devastating impacts for animal health and welfare, food production systems and the economy in Australia. Feral pig populations are capable of spreading and maintaining these diseases in the environment, so it is critical to understand what level of contact occurs within and between feral pig populations to better inform disease preparedness. Here, we developed a proximity-based social network for feral pigs in eastern Australia, which elucidates the contact rate among individual pigs, both within and between sounders. The data procured from this network are critical for refining disease models and enhancing preparedness strategies in Australia. This study further revealed that the contact rate among feral pigs is dependent on two primary factors: the sex of the dyad and the season. It was observed that contacts were more prevalent among female dyads. Additionally, a seasonal pattern was discerned in the contact rate, with summer registering the highest frequency of contact. This research is the first of its kind in Australia, offering new insights into the social behaviour of feral pig populations.

### Network

In our comparison of global network measures between direct and indirect networks, we observed that the global transitivity, average local transitivity, and edge density were marginally higher for the indirect networks. This outcome aligns with our expectations, given that indirect networks permit a greater number of connections. This is due to the inclusion of associations between animals that occur when they share the same space within a window of five days. Understanding this indirect network is crucial for assessing the risk of pathogen dispersal, particularly when the pathogens exhibit prolonged persistence in the environment.

On the other hand, for the node-level networks measures, we found that females had higher strength in the direct network in comparison with males, likely due to greater group cohesion of females within a sounder [19]. In contrast males had a positive effect on betweenness in the indirect netwrok, indicating that they are likely to connect independent groups through individuals with a higher tendency to move between different groups [24]. The role of male feral pigs in connecting sounders should have important implications for the design of disease or population management strategies. For example, removal of adult females is typically seen as more important for sustained control programs aiming to reduce density dependent damage caused by feral pigs because these individuals make the greatest contribution to population growth. However, the removal of adult males may be more important for control programs that aim to contain outbreaks of infectious disease because these individuals are likely to make the greatest contribution to disease transmission between sounders. Intensive population control can also cause surviving animals to change their spatial behaviour, resulting in increased risk of transmission or spillover [25].

### Home range

Previous research that studied four of the feral pig populations included in this study, found that home range was not affected by season [26]. In this study, we did not explore the effect of season on feral pig home range, but rather focused on home range overlap between a dyad. The results of our analyses indicated that dyad (pair of individuals) had a greater home range overlap during summer and followed by autumn.

### Contact rate

The ability for disease to spread relies on many factors. Both direct and indirect contact with pathogens can have significant impact of disease transmission, and so it is important to consider both for epidemiological modelling. Direct contact, where an infected pig is in physical contact with a susceptible pig, is the primary method of transmission for diseases such as ASF and Food Mouth Disease. However, these diseases (e.g. ASF being viable for up to 5 days at 25C° [27]), may also have indirect methods of transmission, for example, through ingestion of contaminated feed, survival of the pathogen in the soil or water sources or possible vectors, such as ticks. Understanding rates of indirect contacts are therefore highly relevant in understanding the risk of disease transmission.

Most **contact occurred within sounders**. We also found that the sex of the dyad statistically affects the contact rate. In our study we found that female-female dyads were most frequent within the same sounder, which can be partly explained by the fact that sounders are composed mainly by adult females, sub-adults and juveniles [19, 28, 29]. Our results differ from a previous study on feral pigs in the United States, which described no significant effect of sex for either direct contact or indirect contact [23]. This could be partly explained by the tendency of females to travel significantly less distance per day, with an average of 3.6 km compared to 4.9 km for males and therefore having more frequent female-female contacts [30].

The estimated contact rate was higher within animals from the same sounder than between sounder, which mean that disease could spread quickly within a sounder and may take longer between sounders. This result has been previously reported for wild boar populations in Europe, where they also found that contact rate between sounders were dependent on the distance between the groups [14]. Moreover, within the sounder, the sex of the dyad played a significant role in the direct contact rate, while between sounder this dyad’s sex is not statistically significant.

For **contact rates between sounders**, we found that sex influenced the rate for indirect contact only, with fewer contacts between males when compare female-female. This results are in contrast to a study on wild boars in Europe that described no effect of animal sex on the association between sounders [14]. For both direct and for indirect contact rates between sounders, male-male dyad and female-male dyad had a negative effect.

We found that the contact rate for within and between sounders was dynamic across the different seasons, which is similar to what has previously been described in wild boars [31]. We found significant differences between seasons with a higher contact rate in summer. This difference in seasonality is critical for understanding management of feral pig populations, but are also important for optimising disease transmission models such as the AADIS model [32]. The incursion and transmission of a new virus into a susceptible feral pig population may spread differently under varying conditions depending on season. For example, controlling an outbreak during summer would potentially require more resources than an outbreak in other season due to the higher number of contacts between individuals during summer. The findings from this study suggest that the incursion of a virus in summer would have a higher impact on the feral pig population (between and within sounders) than comparatively if it were introduced in autumn.

### Limitations

Different populations in our study had varying numbers of collared individuals, with some populations having only two individuals at certain times. This variability in sample size across populations is a limitation when interpreting the results. Small populations are often the result of a few individuals being trapped and collared, and this does not necessarily reflect the actual number of individuals in those groups. This issue creates bias when calculating connections between individuals. For those populations with only a few collared animals, we may observe very few connections and low network measures. However, this could be due to sparse information rather than a true reflection of the population’s structure.

One of the populations (Gebar) is in a small island, which also may incorporate some bias to the network for this location as individuals in that population may have higher contact due to the limited space available.

### Conclusion

Our study highlights the role of sex and seasonality in contact rates, with implications for disease spread and control strategies. Specifically, the higher betweenness of males suggests their crucial role in disease transmission between groups, indicating that removal of adult males may be more effective in containing disease outbreaks. Furthermore, the higher contact rate observed in summer suggests that a virus introduced during this season could have a more significant impact on feral pig populations. These findings have important implications for disease modelling and management of feral pig populations in Australia, aiding in the development of more effective and targeted strategies for disease control and population management.

## Methods

### Ethical approval for the study

Data from feral pigs fitted with GPS-tracking collars were provided by the Queensland Department of Agriculture and Fisheries (DAF) and the New South Wales Department of Primary Industries (DPI). Feral pig GPS-tracking for animals in the state of New South Wales was approved by Animal Research Authority ORA 21-24-003 issued by the NSW Department of Primary Industries Orange Animal Ethics Committee. Animal use for research in Queensland were conducted under approval by the University of New England Animal Ethic Committee (AEC 16-115, AEC 20-023 and AEC 22-056).

### Data sources and data management

We used geolocation data of 74 individual feral pigs from NSW and 72 animals from QLD, for a total of 146 feral pigs collared between 2017 and 2023 for QLD and between 2021 and 2023 for NSW (Figure 1). These animals were from a total of 11 different populations from NSW (n=6) and QLD (n=5). The recording intervals for individuals collared in QLD was 30 minutes and data were cleaned by removing points recorded prior to capture, during the first two days after collaring, post mortality, and any inaccuracies [26]. NSW individuals had one-hour recording intervals and data were filtered the same as QLD individuals with the additional removal of points with incomplete or low-quality fixes, and locations confirmed by less than three satellites [33].

Contacts between individuals can be underestimated when the temporal resolution of GPS data is coarser than 30 min [34], therefore we incorporated continuous time movement models (CTMMs) for both NSW and QLD datasets to fit GPS tracking data to infer trajectories at five-minute intervals [17, 34]. This was done by fitting the GPS data using Ornstein-Uhlenbeck F (OUF) with ctmm.guess() function in the *ctmm* R package (Calabrese et al. 2016), which was then extrapolated in five-minute intervals using the predicted fit model (code adapted from Yang et al. 2023).

### Temporal and spatial thresholds for direct and indirect contact

Direct contact was defined when two individuals interacted either at 2, 5, or 350 metre buffers within a five-minute interval [34]. A previues study used 350 metres as a spatial threshold [14], while other use the approximate average body length of an individual [34]. ASF is estimated to remain infectious in the environment for five days [34] and therefore for the purpose of our study an indirect contact was defined when two individuals interacted within a 2 or 5 metre buffer within a five day interval. These spatiotemporal interaction thresholds aimed to represent indirect contact at populated locations (e.g. waterholes) that are potential hotspots for pathogen exposure. Temporal groups for direct and indirect contact were calculated with R package spatsoc [35] group_times() function using the temporal thresholds listed above [35]. Then spatial thresholds were calculated with spatsoc group_pts() function using a threshold of 0.000045 degrees (∼5 metres) that was split by pig population, year, and season [35]. In the networks, each individual represents a node, and the contact between two individuals represents an edge. A spatial matrix was then created for direct and indirect contact by calculating the edge distances of each individual with the r package spatsoc edge_dist() function using the same degree threshold and split parameters as listed above [35].

### Network analysis

To estimate the contact between individuals, networks were created for each population and year [24, 36]. Proximity-based individual networks were built using the location of each individual in both space and time using the *spatsoc* R package [35]. Spatial-temporal matrices calculated for specific indirect and direct contact thresholds, described above, were used to create separate networks for indirect and direct contact. First, individual matrixes were created from the spatial-temporal matrices with r package spatsoc get_gbi() function, and then a network was built with the get_network() function from the *asnipe* R package using SRI (simple ratio index) as the association index [37]. An undirected weighted network adjacency matrix, which assigns nodes different values depending on the number of associations the connected nodes have, was created for each population in each recorded year with the R package [38].

Networks were visualised with the R package *ggraph* [39]. Node-level network metrics were further calculated with the R package *igraph* [38] including degree centrality, strength, and betweenness for each node. Degree centrality quantifies the number of connections an individual has. The more connections an individual has, the higher their degree centrality. Strength, on the other hand, encapsulates the robustness of these connections. The more time two individuals spend connected, the greater the strength [24]. Lastly, node betweenness measures the number of paths that pass through a node (or individual) to connect two other nodes. In other words, it quantifies how often an individual serves as the quickest link between two other individuals in a network. Betweenness is a crucial measure in the context of disease transmission, as it indicates how frequently an individual could potentially transmit a disease between two others. Global network measures were also calculated, including, average local transitivity, global transitivity, edge density, number of nodes, mean distance, and number of edges for each population and by year. Transitivity measures the likelihood that the adjacent edges of a node are connected (ratio of count of triangles). The local transitivity gives a measure of how interconnected a node’s neighbours, and the global transitivity measure the overall level of transitivity in the network. [40]. We also calculated these measures for the three thresholds (2, 5 and 350m) and compared the differences between the structures of the networks based on the thresholds.

### Home range estimation

Home range is defined as the area utilised by each animal for its normal activities of foraging, mating and caring for young [41]. Average seasonal home ranges for each individual in each recorded season (Summer: December-February; Autumn: March-May; Winter: June-August; Spring: September-November) were estimated for populations with more than three individuals. Home ranges were determined by first calculating the kernel density with 95% confidence with the *amt* R package hr_kde() function [42]. The home range overlap was then determined by comparing the proportions of kernel densities that overlap between each pair of individuals using the amt R package hr_overlap() function [42]. To determine if dyads were comparing individuals within or between sounders, dyads with home range overlaps over 50% were considered to belong within the same sounder, and dyads with home range overlaps under 50% were considered to belong to different sounders [29].

### Contact rates

Direct and indirect contact rates within and between sounders at each site, as well as overall expected contact rate, were estimated for each year, season, and pig population. The contact rate refers to the frequency at which two specific individuals come into contact per unit of time. Contact rates were determined by first calculating how many days each individual was recorded in the direct and indirect spatial and temporal threshold and then determining if the dyad being compared was within or between different sounders, and the type of contact based on sex (female-female, male-male, or female-male). Contact rates were then calculated for each dyad by dividing the number of contact observations from the dyad distance matrix by the minimum number of days for the individual with the least recordings. Mean direct and indirect contact rates and standard deviation were calculated for each type of contact or dyad, which refers to a pair of individuals (female-female, male-male, female-male) between and within sounders for each population for every season and year.

### Statistical analyses

We used three linear mixed effects models to estimate the effects of sex, season, and type of contact (between or within sounders) on direct contact rates. Parameters were selected based on the results of null hypothesis testing. The first model specified overall contact rate (between and within sounders) as the dependent variable and included fixed effects for type of contact (between or within sounder), sex of the dyad, and season. Pig population was included as a random effect. The second model used direct contact rates within sounders as the dependent variable and included the same fixed and random effects as the first model, excluding type of contact. The third model included the same fixed and random effects as the second model, using direct contact rates between sounders as the dependent variable. We then repeated the process using indirect contact rates for the dependent variables. Models were fitted using the lme4 package [43] for R. To determine if sex influences the network measures (degree, centrality, betweenness and strength) we performed a Wilcoxon ran-sum test, using the networks build with five metres threshold and for direct and indirect networks. The code can be found in GitHub.

## Acknowledgements

The authors want to thank Southern Queensland Landscapes for providing data pertaining to this study. We acknowledge the HEAL (Healthy Environments and Lives) National Research Network, which receives funding from the National Health and Medical Research Council (NHMRC) Special Initiative in Human Health and Environmental Change (2008937). We acknowledge NSW Local Land Services and Southern Queensland Landscapes for their help with the trapping and collaring of the animals.

## References

1. Risch, D.R., J. Ringma, and M.R. Price, The global impact of wild pigs (Sus scrofa) on terrestrial biodiversity. Scientific Reports, 2021. 11(1).

2. Pimental, D.R. Environmental and economic costs of vertebrate species invasions into the United States. 2007.

3. !!! INVALID CITATION !!! [3-6].

4. Buetre, B., Potential socio-economic impacts of an outbreak of foot-and-mouth disease in Australia. 2013.

5. Slatyer, R., et al., Potential economic consequences of African swine fever in Australia. 2023.

6. Pearson, H.E., Understanding and mitigating the risk of pathogen transmission from wild animals to domestic pigs in Australia. 2012. p. 252–252.

7. Orr, B., et al., Leptospirosis is an emerging infectious disease of pig-hunting dogs and humans in North Queensland. PLOS Neglected Tropical Diseases, 2022. 16(1): p. e0010100.

8. Cooper, A., et al., Detection of Coxiella burnetii dna in wildlife and ticks in Northern Queensland, Australia. Vector-Borne and Zoonotic Diseases, 2013.

9. Northern Territory Government. Japanese encephalitis in animals. 2022 [cited 2024 24/06/2024]; Available from: https://nt.gov.au/industry/agriculture/livestock-and-animals/animal-health-and-diseases/japanese-encephalitis-in-animals#:∼:text=Northern%20Territory,and%20Cox%2DDaly%20unincorporated%20areas.

10. Bradhurst, R.A., et al., A hybrid modeling approach to simulating foot-and-mouth disease outbreaks in Australian livestock. Frontiers in Environmental Science, 2015. 3: p. 17.

11. Craft, M.E., Infectious disease transmission and contact networks in wildlife and livestock. Philosophical Transactions of the Royal Society B: Biological Sciences, 2015. 370(1669): p. 20140107.

12. Taylor, R.A., et al., The Risk of Infection by African Swine Fever Virus in European Swine Through Boar Movement and Legal Trade of Pigs and Pig Meat. Frontiers in Veterinary Science, 2020. 6.

13. Taylor, R.A., et al., Predicting spread and effective control measures for African swine fever—Should we blame the boars? Transboundary and Emerging Diseases, 2021. 68(2): p. 397–416.

14. Podgórski, T., M. Apollonio, and O. Keuling, Contact rates in wild boar populations: Implications for disease transmission. The Journal of Wildlife Management, 2018. 82(6): p. 1210–1218.

15. Anderson, R.M. and R.M. May, Infectious diseases of humans: Dynamics and control. 1992, Oxford University.

16. Lloyd-Smith, J.O., et al., Superspreading and the effect of individual variation on disease emergence. Nature, 2005. 438(7066): p. 355–359.

17. Calabrese, J.M., C.H. Fleming, and E. Gurarie, ctmm: anR package for analyzing animal relocation data as a continuous-time stochastic process. Methods in Ecology and Evolution, 2016. 7(9): p. 1124–1132.

18. Albery, G.F., et al., Unifying spatial and social network analysis in disease ecology. Journal of Animal Ecology, 2021. 90(1): p. 45–61.

19. Spencer, P.B.S., et al., The sociogenetic structure of a controlled feral pig population. Wildlife Research, 2005. 32(4): p. 297–304.

20. Podgórski, T., et al., Long-Lasting, Kin-Directed Female Interactions in a Spatially Structured Wild Boar Social Network. Plos One, 2014. 9(6).

21. Titus, C.L., et al., Genomic tools reveal complex social organization of an invasive large mammal (Sus scrofa). Biological Invasions, 2022. 24(10): p. 3199–3216.

22. Mayer, J.J., Wild Pigs behavior. Wild pigs: biology, damage, control techniques and management. Savannah River National Laboratory SRNL-RP-2009-00869, Aiken, SC, USA, 2009: p. 221–246.

23. Pepin, K.M., et al., Contact heterogeneities in feral swine: implications for disease management and future research. Ecosphere, 2016. 7(3): p. e01230.

24. Farine, D.R. and H. Whitehead, Constructing, conducting and interpreting animal social network analysis. Journal of Animal Ecology, 2015. 84(5): p. 1144–1163.

25. Ham, C., et al., Effect of culling on individual badger behaviour: Potential implications for bovine tuberculosis transmission. Journal of Applied Ecology, 2019. 56(11): p. 2390–2399.

26. Wilson, C., M. Gentle, and D. Marshall, Factors influencing the activity ranges of feral pigs (Sus scrofa) across four sites in eastern Australia. Wildlife Research, 2023.

27. Mazur-Panasiuk, N. and G. Wozniakowski, Natural inactivation of African swine fever virus in tissues: Influence of temperature and environmental conditions on virus survival. Vet Microbiol, 2020. 242: p. 108609.

28. Boitani, L., et al., Spatial and Activity Patterns of Wild Boars in Tuscany, Italy. Journal of Mammalogy, 1994. 75(3): p. 600–612.

29. Gabor, T.M., et al., Demography, sociospatial behaviour and genetics of feral pigs Sus scrofa in a semi-arid environment. Journal of Zoology, 1999. 247(3): p. 311–322.

30. Wilson, C., M. Gentle, and D. Marshall, Feral pig (Sus scrofa) activity and landscape feature revisitation across four sites in eastern Australia. Australian Mammalogy, 2023.

31. Yang, A., et al., Effects of social structure and management on risk of disease establishment in wild pigs. Journal of Animal Ecology, 2021. 90(4): p. 820–833.

32. Bradhurst, R., et al., Modelling the spread and control of African swine fever in domestic and feral pigs. 2021.

33. Bjørneraas, K., et al., Screening global positioning system location data for errors using animal movement characteristics. The Journal of Wildlife Management, 2010. 74(6): p. 1361–1366.

34. Yang, A., et al., Deriving spatially explicit direct and indirect interaction networks from animal movement data. Ecol Evol, 2023. 13(3): p. e9774.

35. Robitaille, A.L., Q.M.R. Webber, and E.V. Wal, Conducting social network analysis with animal telemetry data: Applications and methods using spatsoc. Methods in Ecology and Evolution, 2019. 10(8): p. 1203–1211.

36. Dougherty, E.R., et al., Going through the motions: incorporating movement analyses into disease research. Ecology Letters, 2018. 21(4): p. 588–604.

37. Farine, D.R., Animal social network inference and permutations for ecologists in R using. Methods in Ecology and Evolution, 2013. 4(12): p. 1187–1194.

38. Gabor Csardi and T. Nepusz, The igraph software package for complex network research. InterJournal, 2006. Complex Systems: p. 1965.

39. Pedersen, T.L., ggraph: An Implementation of Grammar of Graphics for Graphs and Networks. 2022.

40. Csárdi, G. and T. Nepusz, The igraph software package for complex network research. InterJournal Complex Systems, 2006. 1695: p. 1–9.

41. Burt, W.H., Territoriality and Home Range Concepts as Applied to Mammals. Journal of Mammalogy, 1943. 24(3): p. 346–352.

42. Signer, J., J. Fieberg, and T. Avgar, Animal movement tools (amt): R package for managing tracking data and conducting habitat selection analyses. Ecology and evolution, 2019. 9(2): p. 880–890.

43. Bates, D., et al., Fitting Linear Mixed-Effects Models Using lme4. Journal of Statistical Software, 2015. 67(1): p. 1 – 48.

