## Supplementary material for "Quantifying Feral Pig Interactions to Inform Disease Transmission Networks"

***Results***

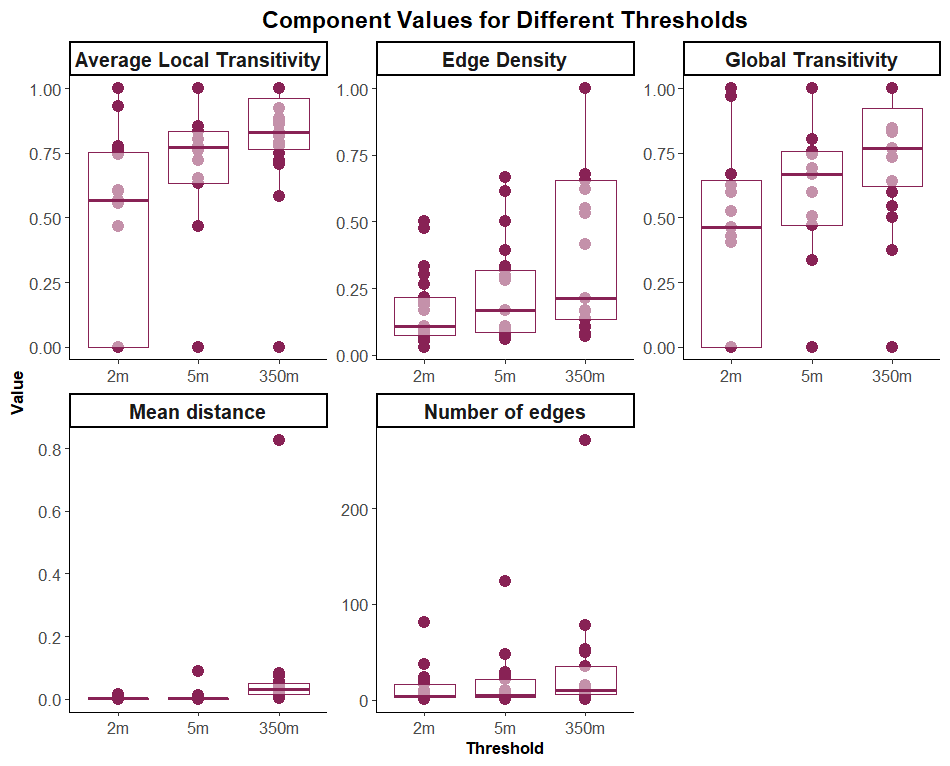

*Figure S1. Sensitivity analysis. Global network measures derived from thresholds of 2, 5, and 350 metres.*

| *a)* | 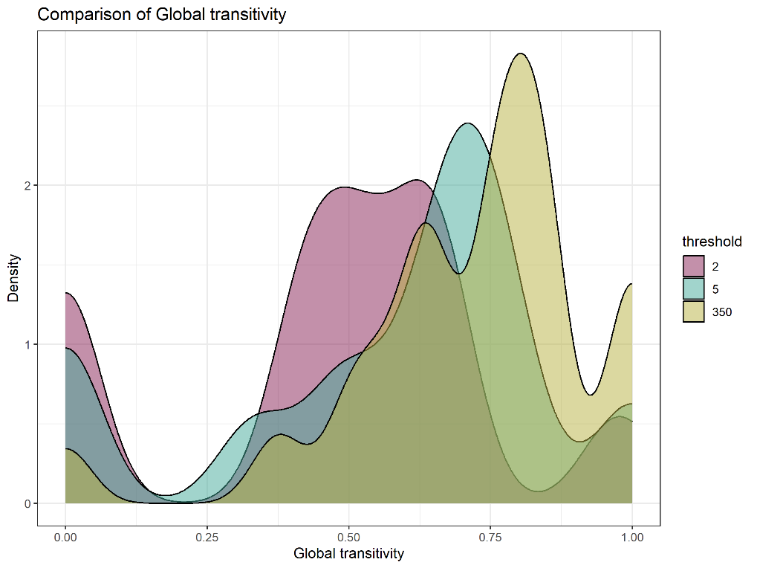 |
| --- | --- |
| *b)* | 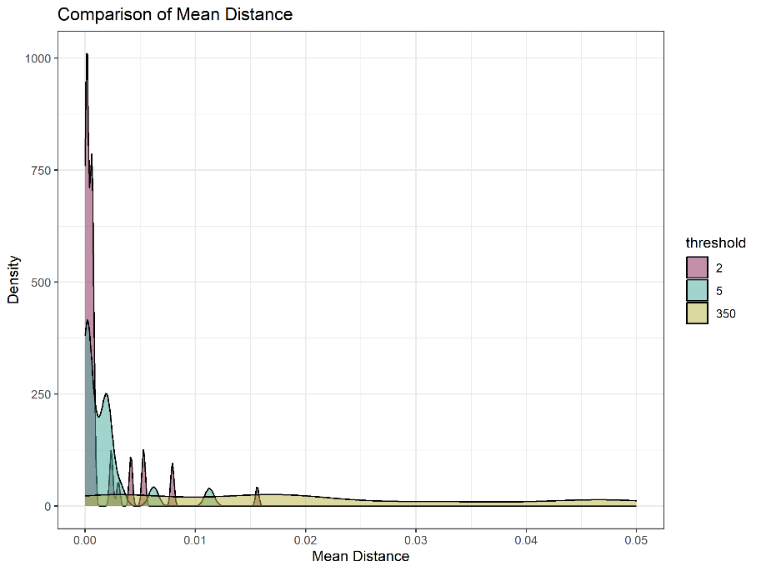 |
| *c)* | 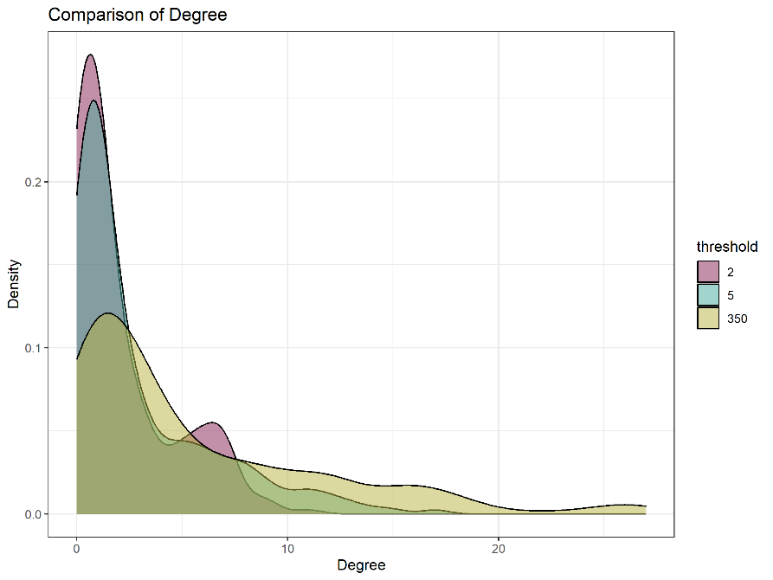 |

*Figure S2. Density comparison between different thresholds. a) global transitivity, b) mean distance, and c) degree*

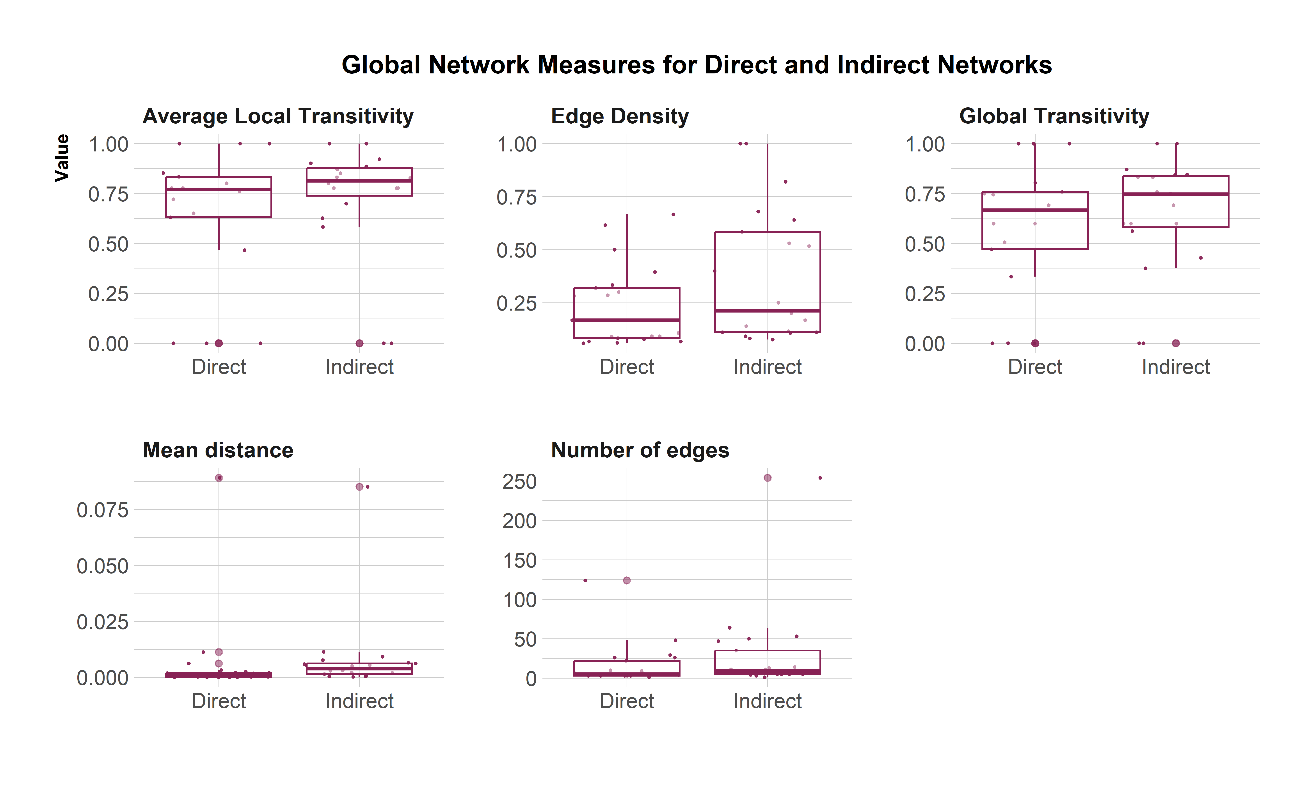

Figure S3. Comparison between global network measures, including average local transitivity, edge density, global transitivity, mean distance and number of edges for direct and indirect networks using a 5 metres threshold.

| *Direct contact 5 metres* | *Indirect contact 5 metres* |
| --- | --- |
| 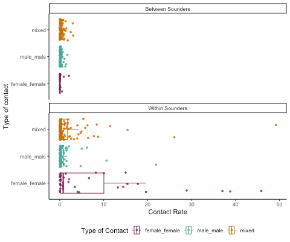 | 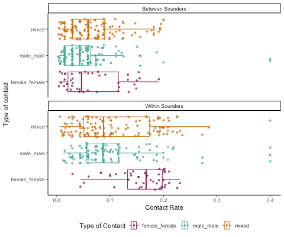 |

*Figure S4. Direct and indirect contacts between and within sounders by type of contact (female-female, male-male and mixed) with a 5 metres threshold*

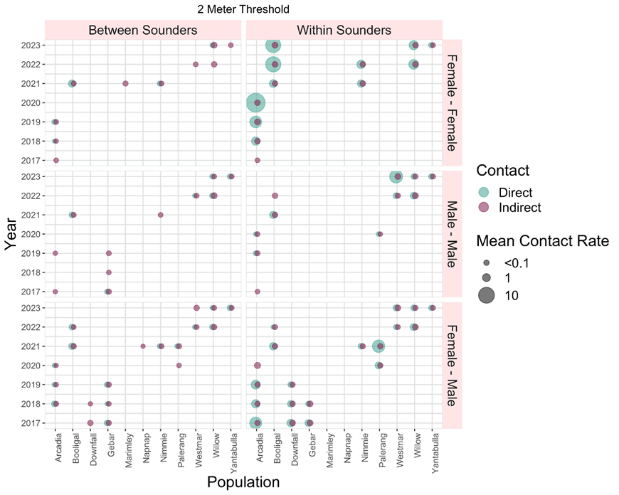

Figure S5. Diagram of direct and indirect mean contact rate for dyads between sounders and within sounders per year and population with a 2 metres threshold

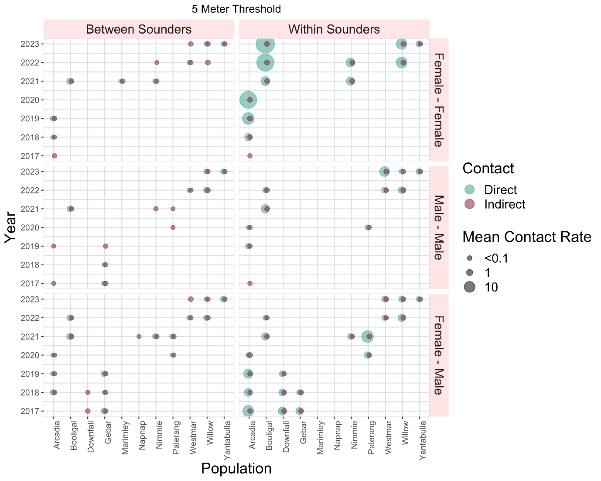

Figure S6. Diagram of direct and indirect mean contact rate for dyads between sounders and within sounders per year and population with a 5 metres threshold

*Table S1. Summary of the mean direct contact rate per population, year, season and type of dyad.*

| *population* | *year* | *season* | *Dyad sex* | *Contact type (1=within and 0=between sounders)* | *Total associations* | *N* | *mean* | *SD* |
| --- | --- | --- | --- | --- | --- | --- | --- | --- |
| *Arcadia* | *2017* | *Autumn* | *mixed* | *0* | *4* | *2* | *2.550649* | *0.128457* |
| *Arcadia* | *2017* | *Autumn* | *mixed* | *1* | *4* | *4* | *99.46101* | *20.75192* |
| *Arcadia* | *2017* | *Spring* | *mixed* | *0* | *6* | *5* | *1.798635* | *0.909897* |
| *Arcadia* | *2017* | *Spring* | *mixed* | *1* | *6* | *4* | *2.053822* | *1.634365* |
| *Arcadia* | *2017* | *Summer* | *mixed* | *0* | *6* | *5* | *2.152549* | *0.863408* |
| *Arcadia* | *2017* | *Summer* | *mixed* | *1* | *6* | *4* | *2.745296* | *1.603521* |
| *Arcadia* | *2017* | *Winter* | *mixed* | *0* | *4* | *2* | *0.964113* | *0.835425* |
| *Arcadia* | *2017* | *Winter* | *mixed* | *1* | *4* | *4* | *51.51758* | *42.08773* |
| *Arcadia* | *2018* | *Autumn* | *female_female* | *0* | *9* | *2* | *0.01099* | *0.000121* |
| *Arcadia* | *2018* | *Autumn* | *female_female* | *1* | *9* | *2* | *4.650671* | *1.542155* |
| *Arcadia* | *2018* | *Autumn* | *mixed* | *0* | *9* | *5* | *0.591523* | *0.371429* |
| *Arcadia* | *2018* | *Autumn* | *mixed* | *1* | *9* | *6* | *3.154798* | *1.914211* |
| *Arcadia* | *2018* | *Spring* | *female_female* | *0* | *9* | *2* | *0.01105* | *8.63E-05* |
| *Arcadia* | *2018* | *Spring* | *female_female* | *1* | *9* | *4* | *4.628381* | *1.565643* |
| *Arcadia* | *2018* | *Spring* | *mixed* | *0* | *9* | *4* | *0.283849* | *0.163809* |
| *Arcadia* | *2018* | *Spring* | *mixed* | *1* | *9* | *4* | *0.476719* | *0.225275* |
| *Arcadia* | *2018* | *Summer* | *female_female* | *0* | *9* | *2* | *0.011111* | *NA* |
| *Arcadia* | *2018* | *Summer* | *female_female* | *1* | *9* | *4* | *9.440249* | *2.478648* |
| *Arcadia* | *2018* | *Summer* | *mixed* | *0* | *9* | *4* | *0.496774* | *0.096774* |
| *Arcadia* | *2018* | *Summer* | *mixed* | *1* | *9* | *4* | *1.172561* | *0.397686* |
| *Arcadia* | *2018* | *Winter* | *female_female* | *0* | *8* | *2* | *0.01099* | *0.000121* |
| *Arcadia* | *2018* | *Winter* | *female_female* | *1* | *8* | *2* | *4.650671* | *1.542155* |
| *Arcadia* | *2018* | *Winter* | *mixed* | *0* | *8* | *5* | *0.506759* | *0.376197* |
| *Arcadia* | *2018* | *Winter* | *mixed* | *1* | *8* | *4* | *0.451905* | *0.205161* |
| *Arcadia* | *2019* | *Autumn* | *female_female* | *0* | *4* | *3* | *0.25726* | *0.098542* |
| *Arcadia* | *2019* | *Autumn* | *mixed* | *0* | *4* | *2* | *0.095222* | *0.059* |
| *Arcadia* | *2019* | *Spring* | *female_female* | *0* | *8* | *4* | *6.520511* | *0.65099* |
| *Arcadia* | *2019* | *Spring* | *female_female* | *1* | *8* | *2* | *0.855315* | *0.017242* |
| *Arcadia* | *2019* | *Spring* | *male_male* | *1* | *8* | *2* | *0.142857* | *0* |
| *Arcadia* | *2019* | *Spring* | *mixed* | *0* | *8* | *2* | *1.115385* | *0* |
| *Arcadia* | *2019* | *Spring* | *mixed* | *1* | *8* | *5* | *5.724787* | *2.670589* |
| *Arcadia* | *2019* | *Summer* | *female_female* | *0* | *11* | *5* | *18.25723* | *4.175272* |
| *Arcadia* | *2019* | *Summer* | *female_female* | *1* | *11* | *2* | *2.496893* | *0.04851* |
| *Arcadia* | *2019* | *Summer* | *male_male* | *1* | *11* | *2* | *0.091082* | *0.030675* |
| *Arcadia* | *2019* | *Summer* | *mixed* | *0* | *11* | *4* | *3.028806* | *2.964659* |
| *Arcadia* | *2019* | *Summer* | *mixed* | *1* | *11* | *3* | *5.554101* | *2.113131* |
| *Arcadia* | *2019* | *Winter* | *female_female* | *0* | *5* | *4* | *10.25005* | *1.45078* |
| *Arcadia* | *2019* | *Winter* | *female_female* | *1* | *5* | *2* | *1.357959* | *0.026383* |
| *Arcadia* | *2019* | *Winter* | *mixed* | *0* | *5* | *2* | *0.491525* | *0* |
| *Arcadia* | *2019* | *Winter* | *mixed* | *1* | *5* | *2* | *2.024473* | *0.961728* |
| *Arcadia* | *2020* | *Autumn* | *female_female* | *1* | *4* | *2* | *16.02645* | *4.841203* |
| *Arcadia* | *2020* | *Autumn* | *male_male* | *1* | *4* | *2* | *0.019941* | *0.008436* |
| *Arcadia* | *2020* | *Autumn* | *mixed* | *0* | *4* | *2* | *0.032788* | *0.000196* |
| *Arcadia* | *2020* | *Autumn* | *mixed* | *1* | *4* | *2* | *0.026488* | *0.005434* |
| *Arcadia* | *2020* | *Spring* | *female_female* | *1* | *4* | *2* | *17.88264* | *5.401279* |
| *Arcadia* | *2020* | *Spring* | *male_male* | *1* | *4* | *2* | *0.020001* | *0.008421* |
| *Arcadia* | *2020* | *Spring* | *mixed* | *0* | *4* | *2* | *0.032967* | *0* |
| *Arcadia* | *2020* | *Spring* | *mixed* | *1* | *4* | *2* | *0.026567* | *0.005351* |
| *Arcadia* | *2020* | *Summer* | *female_female* | *1* | *4* | *2* | *24.43961* | *7.381748* |
| *Arcadia* | *2020* | *Summer* | *male_male* | *1* | *4* | *2* | *0.025* | *0.011785* |
| *Arcadia* | *2020* | *Summer* | *mixed* | *0* | *4* | *2* | *0.05* | *0* |
| *Arcadia* | *2020* | *Summer* | *mixed* | *1* | *4* | *2* | *0.033333* | *0* |
| *Arcadia* | *2020* | *Winter* | *female_female* | *1* | *4* | *2* | *16.02645* | *4.841203* |
| *Arcadia* | *2020* | *Winter* | *male_male* | *1* | *4* | *2* | *0.019941* | *0.008436* |
| *Arcadia* | *2020* | *Winter* | *mixed* | *0* | *4* | *2* | *0.032788* | *0.000196* |
| *Arcadia* | *2020* | *Winter* | *mixed* | *1* | *4* | *2* | *0.026488* | *0.005434* |
| *Downfall* | *2017* | *Spring* | *mixed* | *1* | *4* | *4* | *0.882118* | *0.851185* |
| *Downfall* | *2017* | *Summer* | *mixed* | *1* | *4* | *4* | *1.349816* | *0.744961* |
| *Downfall* | *2017* | *Winter* | *mixed* | *1* | *5* | *5* | *0.967351* | *0.641151* |
| *Downfall* | *2018* | *Autumn* | *mixed* | *1* | *8* | *8* | *1.424763* | *0.593969* |
| *Downfall* | *2018* | *Spring* | *mixed* | *1* | *8* | *8* | *1.429021* | *0.599714* |
| *Downfall* | *2018* | *Summer* | *mixed* | *1* | *8* | *8* | *1.426364* | *0.610519* |
| *Downfall* | *2018* | *Winter* | *mixed* | *1* | *8* | *8* | *1.424763* | *0.593969* |
| *Downfall* | *2019* | *Autumn* | *mixed* | *0* | *6* | *2* | *0.314951* | *0.258708* |
| *Downfall* | *2019* | *Autumn* | *mixed* | *1* | *6* | *6* | *0.796855* | *1.011457* |
| *Downfall* | *2019* | *Spring* | *mixed* | *0* | *6* | *2* | *0.315756* | *0.258022* |
| *Downfall* | *2019* | *Spring* | *mixed* | *1* | *6* | *6* | *0.925028* | *1.05183* |
| *Downfall* | *2019* | *Summer* | *mixed* | *0* | *6* | *2* | *0.418079* | *0.262034* |
| *Downfall* | *2019* | *Summer* | *mixed* | *1* | *6* | *6* | *1.101194* | *1.178271* |
| *Downfall* | *2019* | *Winter* | *mixed* | *0* | *6* | *2* | *0.314951* | *0.258708* |
| *Downfall* | *2019* | *Winter* | *mixed* | *1* | *6* | *6* | *0.796855* | *1.011457* |
| *Gebar* | *2017* | *Spring* | *male_male* | *0* | *3* | *2* | *0.051392* | *0.01615* |
| *Gebar* | *2017* | *Spring* | *mixed* | *0* | *3* | *2* | *0.339184* | *0.016023* |
| *Gebar* | *2017* | *Spring* | *mixed* | *1* | *3* | *2* | *1.017552* | *0.047324* |
| *Gebar* | *2017* | *Summer* | *male_male* | *0* | *3* | *2* | *0.053763* | *0.018624* |
| *Gebar* | *2017* | *Summer* | *mixed* | *0* | *3* | *2* | *0.354839* | *0* |
| *Gebar* | *2017* | *Summer* | *mixed* | *1* | *3* | *2* | *1.064516* | *0* |
| *Gebar* | *2018* | *Autumn* | *mixed* | *0* | *3* | *2* | *0.186828* | *0.001693* |
| *Gebar* | *2018* | *Autumn* | *mixed* | *1* | *3* | *3* | *0.132978* | *0.056078* |
| *Gebar* | *2018* | *Spring* | *mixed* | *0* | *3* | *2* | *0.187851* | *0.001053* |
| *Gebar* | *2018* | *Spring* | *mixed* | *1* | *3* | *3* | *0.133706* | *0.056616* |
| *Gebar* | *2018* | *Summer* | *mixed* | *0* | *3* | *2* | *0.188889* | *0* |
| *Gebar* | *2018* | *Summer* | *mixed* | *1* | *3* | *3* | *0.134444* | *0.057667* |
| *Gebar* | *2018* | *Winter* | *mixed* | *0* | *3* | *2* | *0.186828* | *0.001693* |
| *Gebar* | *2018* | *Winter* | *mixed* | *1* | *3* | *3* | *0.132978* | *0.056078* |
| *Gebar* | *2019* | *Autumn* | *mixed* | *0* | *2* | *2* | *0.152437* | *0.070222* |
| *Gebar* | *2019* | *Summer* | *mixed* | *0* | *2* | *2* | *0.174237* | *0.072822* |
| *Gebar* | *2019* | *Winter* | *mixed* | *0* | *2* | *2* | *0.153264* | *0.069971* |
| *Palerang* | *2020* | *Autumn* | *mixed* | *0* | *4* | *2* | *1.095238* | *0.281352* |
| *Palerang* | *2020* | *Autumn* | *mixed* | *1* | *4* | *4* | *10.64286* | *3.549313* |
| *Palerang* | *2020* | *Spring* | *male_male* | *1* | *8* | *2* | *0.136154* | *0.105445* |
| *Palerang* | *2020* | *Spring* | *mixed* | *0* | *8* | *2* | *0.342844* | *0.283801* |
| *Palerang* | *2020* | *Spring* | *mixed* | *1* | *8* | *6* | *3.893684* | *4.055169* |
| *Palerang* | *2020* | *Summer* | *male_male* | *1* | *8* | *2* | *0.184751* | *0.096373* |
| *Palerang* | *2020* | *Summer* | *mixed* | *0* | *8* | *2* | *0.472141* | *0.264589* |
| *Palerang* | *2020* | *Summer* | *mixed* | *1* | *8* | *6* | *5.996972* | *4.433133* |
| *Palerang* | *2020* | *Winter* | *mixed* | *0* | *6* | *2* | *0.277966* | *0.270152* |
| *Palerang* | *2020* | *Winter* | *mixed* | *1* | *6* | *6* | *3.640875* | *3.94511* |
| *Palerang* | *2021* | *Autumn* | *mixed* | *0* | *6* | *2* | *0.083374* | *0.062967* |
| *Palerang* | *2021* | *Autumn* | *mixed* | *1* | *6* | *6* | *10.98369* | *4.089579* |
| *Palerang* | *2021* | *Summer* | *mixed* | *0* | *6* | *2* | *0.108757* | *0.063437* |
| *Palerang* | *2021* | *Summer* | *mixed* | *1* | *6* | *6* | *12.69962* | *3.73483* |
| *Palerang* | *2021* | *Winter* | *mixed* | *0* | *6* | *2* | *0.154762* | *0.013041* |
| *Palerang* | *2021* | *Winter* | *mixed* | *1* | *6* | *6* | *57.71343* | *16.75624* |
| *Westmar* | *2022* | *Spring* | *male_male* | *0* | *11* | *5* | *0.031523* | *0.018466* |
| *Westmar* | *2022* | *Spring* | *male_male* | *1* | *11* | *7* | *1.411675* | *0.855507* |
| *Westmar* | *2022* | *Spring* | *mixed* | *0* | *11* | *5* | *0.091759* | *0.065357* |
| *Westmar* | *2022* | *Spring* | *mixed* | *1* | *11* | *7* | *1.136626* | *0.607836* |
| *Westmar* | *2022* | *Summer* | *male_male* | *0* | *10* | *5* | *0.050691* | *0.017243* |
| *Westmar* | *2022* | *Summer* | *male_male* | *1* | *10* | *7* | *2.323983* | *0.833109* |
| *Westmar* | *2022* | *Summer* | *mixed* | *0* | *10* | *5* | *0.153226* | *0.074663* |
| *Westmar* | *2022* | *Summer* | *mixed* | *1* | *10* | *5* | *1.751083* | *0.544255* |
| *Westmar* | *2022* | *Winter* | *male_male* | *0* | *11* | *5* | *0.036155* | *0.017884* |
| *Westmar* | *2022* | *Winter* | *male_male* | *1* | *11* | *7* | *1.622054* | *0.882943* |
| *Westmar* | *2022* | *Winter* | *mixed* | *0* | *11* | *5* | *0.111539* | *0.07068* |
| *Westmar* | *2022* | *Winter* | *mixed* | *1* | *11* | *7* | *1.232781* | *0.540481* |
| *Westmar* | *2023* | *Summer* | *male_male* | *1* | *4* | *2* | *5.727273* | *0* |
| *Westmar* | *2023* | *Summer* | *mixed* | *1* | *4* | *2* | *0.272727* | *0* |
| *booligal* | *2021* | *Spring* | *female_female* | *0* | *10* | *2* | *0.027757* | *0.006366* |
| *booligal* | *2021* | *Spring* | *female_female* | *1* | *10* | *5* | *4.148873* | *1.949168* |
| *booligal* | *2021* | *Spring* | *male_male* | *0* | *10* | *2* | *0.337049* | *0.104545* |
| *booligal* | *2021* | *Spring* | *male_male* | *1* | *10* | *3* | *2.755503* | *1.077664* |
| *booligal* | *2021* | *Spring* | *mixed* | *0* | *10* | *4* | *0.341175* | *0.142787* |
| *booligal* | *2021* | *Spring* | *mixed* | *1* | *10* | *9* | *3.14066* | *4.520546* |
| *booligal* | *2021* | *Summer* | *female_female* | *0* | *10* | *2* | *0.032258* | *NA* |
| *booligal* | *2021* | *Summer* | *female_female* | *1* | *10* | *5* | *5.960543* | *2.170829* |
| *booligal* | *2021* | *Summer* | *male_male* | *0* | *10* | *2* | *0.391705* | *0.103456* |
| *booligal* | *2021* | *Summer* | *male_male* | *1* | *10* | *3* | *3.192367* | *1.109778* |
| *booligal* | *2021* | *Summer* | *mixed* | *0* | *10* | *2* | *0.258065* | *0* |
| *booligal* | *2021* | *Summer* | *mixed* | *1* | *10* | *8* | *4.290045* | *5.480422* |
| *booligal* | *2022* | *Autumn* | *female_female* | *1* | *5* | *2* | *14.18582* | *4.893425* |
| *booligal* | *2022* | *Autumn* | *mixed* | *0* | *5* | *2* | *0.2* | *0* |
| *booligal* | *2022* | *Autumn* | *mixed* | *1* | *5* | *2* | *0.03297* | *0.000314* |
| *booligal* | *2022* | *Spring* | *female_female* | *1* | *5* | *2* | *14.26348* | *4.919184* |
| *booligal* | *2022* | *Spring* | *mixed* | *0* | *5* | *2* | *0.2* | *0* |
| *booligal* | *2022* | *Spring* | *mixed* | *1* | *5* | *2* | *0.03315* | *0.000201* |
| *booligal* | *2022* | *Summer* | *female_female* | *1* | *5* | *2* | *14.34229* | *4.94603* |
| *booligal* | *2022* | *Summer* | *mixed* | *0* | *5* | *2* | *0.2* | *0* |
| *booligal* | *2022* | *Summer* | *mixed* | *1* | *5* | *2* | *0.033333* | *0* |
| *booligal* | *2022* | *Winter* | *female_female* | *1* | *5* | *2* | *14.18582* | *4.893425* |
| *booligal* | *2022* | *Winter* | *mixed* | *0* | *5* | *2* | *0.2* | *0* |
| *booligal* | *2022* | *Winter* | *mixed* | *1* | *5* | *2* | *0.03297* | *0.000314* |
| *booligal* | *2023* | *Summer* | *female_female* | *1* | *2* | *2* | *8.6* | *0* |
| *nimmie* | *2021* | *Autumn* | *female_female* | *0* | *13* | *6* | *0.277785* | *0.110793* |
| *nimmie* | *2021* | *Autumn* | *female_female* | *1* | *13* | *11* | *7.445236* | *5.617239* |
| *nimmie* | *2021* | *Autumn* | *mixed* | *0* | *13* | *4* | *0.034041* | *0.020222* |
| *nimmie* | *2021* | *Autumn* | *mixed* | *1* | *13* | *10* | *0.355684* | *0.304409* |
| *nimmie* | *2021* | *Spring* | *female_female* | *0* | *11* | *2* | *0.675676* | *0* |
| *nimmie* | *2021* | *Spring* | *female_female* | *1* | *11* | *10* | *7.365741* | *5.253616* |
| *nimmie* | *2021* | *Spring* | *mixed* | *0* | *11* | *3* | *0.031849* | *0.019574* |
| *nimmie* | *2021* | *Spring* | *mixed* | *1* | *11* | *6* | *0.327846* | *0.339278* |
| *nimmie* | *2021* | *Summer* | *female_female* | *1* | *8* | *8* | *13.06781* | *5.547677* |
| *nimmie* | *2021* | *Winter* | *female_female* | *0* | *13* | *6* | *0.286986* | *0.059037* |
| *nimmie* | *2021* | *Winter* | *female_female* | *1* | *13* | *11* | *7.130967* | *5.592559* |
| *nimmie* | *2021* | *Winter* | *mixed* | *0* | *13* | *4* | *0.027317* | *0.01719* |
| *nimmie* | *2021* | *Winter* | *mixed* | *1* | *13* | *10* | *0.304058* | *0.293011* |
| *nimmie* | *2022* | *Autumn* | *female_female* | *0* | *3* | *2* | *0.025146* | *0.019698* |
| *nimmie* | *2022* | *Autumn* | *female_female* | *1* | *3* | *2* | *4.800498* | *3.414369* |
| *nimmie* | *2022* | *Summer* | *female_female* | *0* | *3* | *2* | *0.032284* | *0.021687* |
| *nimmie* | *2022* | *Summer* | *female_female* | *1* | *3* | *2* | *6.163216* | *3.430554* |
| *nimmie* | *2022* | *Winter* | *female_female* | *0* | *3* | *2* | *0.047619* | *NA* |
| *nimmie* | *2022* | *Winter* | *female_female* | *1* | *3* | *2* | *8.522572* | *2.306531* |
| *willow* | *2022* | *Spring* | *male_male* | *0* | *11* | *4* | *3* | *1.290994* |
| *willow* | *2022* | *Spring* | *male_male* | *1* | *11* | *7* | *34.28571* | *16.06885* |
| *willow* | *2022* | *Spring* | *mixed* | *0* | *11* | *4* | *4.142857* | *1.46385* |
| *willow* | *2022* | *Spring* | *mixed* | *1* | *11* | *7* | *41.0303* | *16.32725* |
| *willow* | *2022* | *Summer* | *female_female* | *1* | *13* | *4* | *2.523704* | *1.264902* |
| *willow* | *2022* | *Summer* | *male_male* | *0* | *13* | *4* | *1.469677* | *1.7062* |
| *willow* | *2022* | *Summer* | *male_male* | *1* | *13* | *7* | *18.86937* | *20.70438* |
| *willow* | *2022* | *Summer* | *mixed* | *0* | *13* | *4* | *0.134255* | *0.046172* |
| *willow* | *2022* | *Summer* | *mixed* | *1* | *13* | *8* | *1.310649* | *0.527267* |
| *willow* | *2023* | *Autumn* | *female_female* | *0* | *12* | *2* | *0.222222* | *0* |
| *willow* | *2023* | *Autumn* | *female_female* | *1* | *12* | *2* | *1.855583* | *0.266336* |
| *willow* | *2023* | *Autumn* | *male_male* | *0* | *12* | *2* | *0.024562* | *0.008928* |
| *willow* | *2023* | *Autumn* | *male_male* | *1* | *12* | *6* | *0.592031* | *0.263716* |
| *willow* | *2023* | *Autumn* | *mixed* | *0* | *12* | *5* | *0.206997* | *0.090993* |
| *willow* | *2023* | *Autumn* | *mixed* | *1* | *12* | *7* | *1.606501* | *0.700692* |
| *willow* | *2023* | *Spring* | *female_female* | *1* | *8* | *2* | *1.855583* | *0.266336* |
| *willow* | *2023* | *Spring* | *male_male* | *0* | *8* | *2* | *0.045045* | *0.015604* |
| *willow* | *2023* | *Spring* | *male_male* | *1* | *8* | *2* | *0.767626* | *0.271656* |
| *willow* | *2023* | *Spring* | *mixed* | *0* | *8* | *5* | *0.369079* | *0.207746* |
| *willow* | *2023* | *Spring* | *mixed* | *1* | *8* | *4* | *0.025231* | *0.008374* |
| *willow* | *2023* | *Summer* | *female_female* | *0* | *12* | *2* | *0.033898* | *0* |
| *willow* | *2023* | *Summer* | *female_female* | *1* | *12* | *2* | *2.112926* | *0.076375* |
| *willow* | *2023* | *Summer* | *male_male* | *0* | *12* | *2* | *0.028249* | *0.009786* |
| *willow* | *2023* | *Summer* | *male_male* | *1* | *12* | *6* | *0.742203* | *0.311823* |
| *willow* | *2023* | *Summer* | *mixed* | *0* | *12* | *5* | *0.263923* | *0.103316* |
| *willow* | *2023* | *Summer* | *mixed* | *1* | *12* | *8* | *1.631776* | *0.734409* |
| *willow* | *2023* | *Winter* | *female_female* | *1* | *9* | *2* | *1.855583* | *0.266336* |
| *willow* | *2023* | *Winter* | *male_male* | *0* | *9* | *2* | *0.024562* | *0.008928* |
| *willow* | *2023* | *Winter* | *male_male* | *1* | *9* | *4* | *0.604859* | *0.253632* |
| *willow* | *2023* | *Winter* | *mixed* | *0* | *9* | *5* | *0.206997* | *0.090993* |
| *willow* | *2023* | *Winter* | *mixed* | *1* | *9* | *5* | *0.305827* | *0.342275* |
| *yantabulla* | *2023* | *Spring* | *female_female* | *1* | *26* | *4* | *0.124055* | *0.030672* |
| *yantabulla* | *2023* | *Spring* | *male_male* | *0* | *26* | *17* | *0.282215* | *0.240951* |
| *yantabulla* | *2023* | *Spring* | *male_male* | *1* | *26* | *17* | *0.254837* | *0.154186* |
| *yantabulla* | *2023* | *Spring* | *mixed* | *0* | *26* | *18* | *0.341267* | *0.197191* |
| *yantabulla* | *2023* | *Spring* | *mixed* | *1* | *26* | *15* | *0.250383* | *0.127033* |
